# MU-BRAIN: MUltiethnic Brain Rna-seq for Alzheimer INitiative

**DOI:** 10.1101/2024.02.20.581250

**Authors:** Zikun Yang, Basilio Cieza, Dolly Reyes-Dumeyer, Annie Lee, Yiyi Ma, Elanur Yilmaz, Rafael Lantigua, Gary W Miller, Lewis M Brown, Lawrence Honig, Benjamin Ciener, Sandra Leskinen, Sharanya Sivakumar, Badri Vardarajan, Brittany N Dugger, Lee-Way Jin, Melissa E Murray, Dennis W Dickson, Robert A Rissman, Annie Hiniker, Margaret Pericak-Vance, Jeff Vance, Tatiana M Foroud, Caghan Kizil, Andrew F Teich, Richard Mayeux, Giuseppe Tosto

## Abstract

Alzheimer’s Disease (AD) exhibits a complex molecular and phenotypic profile. Investigating gene expression plays a crucial role in unraveling the disease’s etiology and progression. Transcriptome data across ethnic groups lack, negatively impacting equity in intervention and research.

We employed 565 brains across six U.S. brain banks (*n=*399 non-Hispanic Whites, *n*=113 Hispanics, *n=*12 African Americans) to generated bulk RNA sequencing from prefrontal cortex. We sought to identify cross-ancestry and ancestry-specific differentially expressed genes (DEG) across Braak stages, adjusting for sex, age at death, and RNA quality metrics. We further validated our findings using the Religious Orders Study/Memory Aging Project brains (ROS/MAP; *n=*1,095) and performed metanalysis (*n=*1,660). We conducted Gene Set and Variation and Enrichment analysis (GSVA; GSEA). We employed a machine-learning approach for phenotype prediction and gene prioritization to construct a polytranscriptomics risk score (PTRS) splitting our sample into training and testing sub-samples, either randomly or by ethnicity (“ancestry-agnostic” and “ancestry-aware”, respectively). Lastly, we validated top DEG using single-nucleus RNA sequencing (snRNAseq) data.

We identified several DEG associated with Braak staging: AD-known genes *VGF* (*P_adj_* =3.78E- 07) and *ADAMTS2* (*P_ad j_*=1.21E-04) were consistently differentially expressed across statistical models, ethnicities, and replicated in ROS/MAP. Genes from the heat shock protein (*HSP*) family, e.g. *HSPB7* (*P_adj_* =3.78E-07), were the top differentially expressed genes and replicated in ROS/MAP. Ethnic-stratified analyses prioritized *TNFSF14* and *SPOCD1* as top Hispanics DEG. GSEA highlighted “*Alzheimer disease*” (*P_adj_* =4.24E-06) and “*TYROBP causal network in microglia*” (*P_adj_* =1.68E-08) pathways. Up- and down-regulated genes were enriched in several pathways (e.g. “*Immune response activation signal pathways”*, “*Vesicle-mediated transport in synapse*”, “*cognition*”). Ancestry-agnostic and ancestry-aware PTRS effectively classified brains (AUC=0.77 and 0.73 respectively) and replicated in ROS/MAP. snRNAseq validated prioritized genes, including *VGF (*downregulated in neurons; *P_adj_*=1.1 E-07).

This is the largest diverse AD transcriptome in post-mortem brain tissue, to our knowledge. We identified perturbated genes, pathways and network expressions in AD brains resulting in cross-ethnic and ethnic-specific findings, ultimately highlighting the diversity within AD pathogenesis. The latter underscores the need for an integrative and personalized approach in AD studies.

## Introduction

Alzheimer disease exhibits complexity through the interplay of numerous genetic, epigenetic, and environmental factors contributing to its onset and progression. Brains of individulas with a diagnosis of Alzheimer disease undergo notable neuropathological alterations, including a progressive atrophy, in particular of the hippocampal and cortical regions^1^. Numerous past investigations have led to the discovery of specific genes and proteins integral to the neural system, and several risk factors that influence the onset of Alzheimer disease ^2^, including alternative regulation of gene expression mechanisms, such as mRNA-transcription factor interactions, non-coding RNAs, alternative splicing, and/or copy number variants^3^. Differential gene expression analyses contrasting affected vs. non-affected brains are a primary interest of investigation aiming to uncover pathological mechanisms in Alzheimer disease and other neurological diseases^4^.

Although genetic and environmental factors exhibit variability across different ancestral and ethnic groups^5^, most genomic research, especially in the context of neurodegenerative diseases, remains primarily focused on non-Hispanic White (NHW) individuals. This highlights an underlying health disparity in Alzheimer disease research resulting in a significant gap in understanding disease mechanisms. There is a pressing need to diversify research efforts and extend comprehensive genomic studies to include a broader spectrum of ethnic and ancestral groups^6–8^. In this study, we assembled a large dataset of diverse ethnic backgrounds and detailed neuropathological assessment. In addition to NHW samples, our study includes 113 brains from Hispanic individuals, marking this the largest collection of its kind to date. This dataset allows for a detailed exploration into ethnic-specific molecular profiles and their complex associations with AD.

Braak staging, which refers to the extent and location of hyperphosphorylated tau pathology in the brain, effectively categorizes the progression of AD ^1^. We employed Braak staging as a primary outcome because most of our samples (>90%) were ascertained from individuals with at least some degree of AD pathology, thus lacking true pathology-free control brains. This imbalance necessitated an alternative method of categorization beyond the traditional case vs. control contrast. The strong association between Braak staging and clinical manifestations, i.e. memory deficits or poor clinical dementia ratings, has been extensively studied in previous investigations^9,10^. This methodological choice ensures a thorough and detailed investigation, accommodating the sample composition and enhancing the depth of our insights into the complex nature within the disease’s spectrum. Furthermore, categorization of individuals with and without AD often oversees secondary pathological findings or clumps together those (rare) pathology-free brains with others still exhibiting low-to-medium pathology burden.

We present **MU-BRAIN** (**MU**ltiethnic **B**rain **R**na-seq for **A**lzheimer **IN**itiative), a multiethnic transcriptome: through differential gene expression analysis, ML-based prediction scores, and pathway analyses, we report a complex molecular interplay aiming to foster a more inclusive and representative outlook in AD research.

## Materials and methods

### Description of cohorts and sample filters

We obtained a total of 925 prefrontal cortical brain samples across six distinct U.S. cohorts: New York Brain Bank at Columbia University, University of California Davis, University of California- San Diego, Mayo Clinic Florida, and University of Miami. Additionally, we accessed the data related to the brain samples through the National Comprehensive Repository for Alzheimer’s Disease (NCRAD). Age, sex, self-reported ethnic group, neuropathological diagnoses, RIN (RNA integrity number representing integrity values to RNA measurements) and Postmortem Interval (PMI) were collected. After excluding samples with missing neuropathological diagnosis, primary diagnosis other than Alzheimer’s Disease ^11^, or minimum set of covariate data (sex, age at death, RIN), we retain 565 brains (“*Main model*”). In secondary analyses (detailed in the supplementary material) we additionally included PMI (“*Model 2”*, final *n=*272), which was missing in a significant number of brains. Additionally, we expanded the analyses to all brains with an available Braak stage, regardless of their primary neuropathological diagnosis (i.e. AD + non-AD dementias, with or without PMI available [*n=*720 “*Model 3*” and *n=*398 “*Model 4*”, respectively]).

### RNA Sequencing (RNA-Seq)

Samples were submitted to the New York Genome Center for transcriptome library construction. Total RNA was extracted using Qiagen’s RNeasy Mini Kit, according to manufacturer recommendations. Samples were run on a Bioanalyzer or Fragment Analyzer (Agilent) and quantified using Qubit and Ribogreen. Samples were prepped using Kapa’s Kapa Hyper with RiboErase prep, with a standard input of 100ng, and using unique Illumina barcodes. Libraries were sequenced on a NovaSeq 6000 flow cell using 2x100bp cycles, targeting 60 million reads per sample. The RiboErase library prep avoids 3’ bias found with oligo-dT priming with fragmented RNA, and is ideal for autopsy tissue, allowing us to perform RNA-seq with RINs as low as 4 that results in high-quality FASTQ files that pass QC using *FastQC* v 0.11.8. We found that samples that have RIN values between 2.5-3.9 gave high-quality FASTQ data using the RiboErase prep after passing visual inspection of trace degradation and ribosomal subunit presence (for our project, samples that had medium/low trace degradation and visually distinct ribosomal subunits were sequenced).

Gene counts for each BAM file were calculated using the function *featureCounts*. The integration of several genomic datasets offered the potential to increase statistical power, but this advantage may be compromised by batch effects — unintended data variations arising from technical discrepancies across different batches. To mitigate these batch effects across the six cohorts, we first employed a batch correction method, *ComBat-seq*^12^, to remove batch effect for better statistical power and control of false positives in differential expression analysis. A total of 62,649 genes were initially quantified, but genes exhibiting zero expression across all participants were excluded from subsequent analyses, reducing the dataset to 58,942 informative genes.

### Outcome definition

Considering Braak staging (ordered categorical variable ranging from 0 to VI) could lead to an excessive number of contrasts and a subsequent loss of statistical power, we utilized the R (version 4.2.2) package "*PResiduals*"(version 1.0) to transform the Braak stage into a continuous scale^13^ by converting ordered categorical data into probability-scale residuals, effectively mapping the Braak stage into a continuous interval of (-1,1). Alternatively, we constructed a binary phenotype: late-stage Braak vs. early-stage Braak ( Braak > IV vs. Braak ≤ IV, respectively).

### Differential gene expression (DGE) analysis

The *DESeq2* package(version 1.38.3) was used to identify differentially expressed genes (DEG) associated with the continuous/binary Braak^14^ as previously defined. The DESeq2 ‘*des’* function was used to model the counts following a negative binomial distribution, with mean parameter associated with the covariates included in the statistical models. Due to the distributional assumption on the counts, ‘*des’* function was only capable of processing the integral counts without normalization. As explained previously, we included in the main analyses only brains with a primary neuropathological diagnosis of AD and pathology-free control brains. The main statistical model was defined as: RNA count ∼ Age + Sex + RIN + Braak_continuous_. For Model 2 (RNA count ∼ Age + Sex + RIN + PMI + Ethnicity + Study site + Braak_continuous_) we retained only NHW and Hispanic brain samples, due to the limited representation of African American brains.

The function ‘*plotPCA*’ in *DESeq2* was used to compute the principal components of the gene expression counts.

### Validation dataset: ROS/MAP

We replicated our findings using results from the ROS/MAP database^15^. This external validation contrasted AD pathologically diagnosed brains with AD-free control brains. For consistency, we applied the same criteria to define DEG. To ensure consistency between statistical models implemented between MU-BRAIN and ROS/MAP, the Main model employed RNA quality metrics that captured the distribution of the bases within the transcripts (denoted as “Main model + RNAmetrics”). To do so, we used the function *CollectRnaSeqMetrics* from the *Picard* software^16^ to estimate several parameters as reported in **Supplementary methods**. We further applied “Main model + RNAmetrics” to the ROS/MAP dataset using the *DESeq2* package following an identical pipeline described above to obtain DEG associated with Braak staging.

Finally, we conducted a meta-analysis using *Metal*^17^ across MU-BRAIN and ROS/MAP.

### Stratified analyses by ethnic group

To investigate potential differences in transcriptomic signatures between Hispanics and NHWs, we performed stratified analysis within NHW and Hispanics separately. Furthermore, we tested an additional model by incorporating an interaction term (Braak*ethnicity) within the Main model.

### Gene set variation analysis and functional enrichment analysis

We conducted gene set variation analysis (GSVA) using the *GSVA* package^18^(version 1.36.0) to pinpoint differentially expressed gene ontology (GO) sets. This approach groups biologically related genes into sets. The statistical model and covariates used here were consistent with those applied in the DEG analyses. We employed “Gene Ontology gene sets” from the “Molecular signatures database” (MSigDB)^19^ and conducted the differential expression analysis at the pathway level using the “*limma*” package^20^ (version 3.54.2) because GSVA enrichment scores are continuous.

We performed functional enrichment analysis, i.e. enrichment analysis and Gene Set Enrichment Analysis (GSEA) against GO and WikiPathways databases respectively, using the *clusterProfile* package^21^(version 4.10.0). We conducted GSEA for Wikipathways using the *gseWP* function. We applied the *enrichGO* function to conduct GO enrichment analysis with genes stratified into up- and down-regulated expressions. The GO enrichment analysis performs Fisher’s exact test to identify the overrepresented pathways associated with sets of differentially expressed genes that were previously revealed in the differential gene expression analysis, whereas GSEA assesses pathways for statistical overrepresentation across the entire spectrum of ranked DEG through a permutation test. GSEA and GO enrichment analysis were carried out using DEG from the Main model.

### Multiple testing correction

We applied the Benjamini-Hochberg false discovery rate (FDR) to account for multiple testing across genes and gene-sets, implemented in the *DESeq2*, *limma*, GSVA, and *clusterProfile* packages. We define a significant DEG if FDR_p-value_ < 0.05 and an absolute log_2_-fold change (LFC) threshold >0.15 in differential gene expression analyses, GSVA, and GO enrichment analysis.

### Phenotype prediction using machine learning algorithm

We applied Python (version 3.9.13) library *Scikit-learn* (version 1.0.2) to employ the Random Forest algorithm for the machine learning (ML) approach. More details in the **Supplementary methods**. Random Forest generates a multitude of decision trees, each trained on a different subset of the data and with random feature subsets. The data was partitioned into training and testing sets: 75% of the sample was allocated randomly to the training subset, while the remaining 25% was designated as the testing set. To ensure a balanced representation of samples, the stratification was based on the binary Braak outcome as the stratifying variable.

We employed two distinct models: an “Ethnicity-Agnostic” Model (randomly assigning samples to training and testing samples independently of their ethnicity) and a “Ethnicity-Aware” Model (NHW samples were included in the training subset, while Hispanics were included in the testing subset). We employed a bootstrapping approach to the training and testing data independently. (A) For the training dataset, in the “Ethnicity-Agnostic” Model, we generated 1,000 samples with replacement. Each of these samples consisted of 500 individuals (250 brains with high Braak with replacement from a training subset of 346 brains and 250 brains with low Braak, sampled with replacement from a training subset of 63 brains). (B) The same method was applied to the testing dataset: we generated 1,000 samples sets, each consisting of 126 brains (63 high Braak drawn with replacement from a testing subset of 149 brains, and 63 low Braak brains sampled with replacement from a training subset of 27 brains). We used the 1,000 training datasets to independently train the 1,000 random forest algorithms. Then, we tested the performance of the algorithms using 1,000 testing samples and the average result was reported. The same bootstrapping approach was implemented in the “Ethnicity-Aware” model with the only difference that all brains from the training data were sampled only from NHW, while all brains from the testing data were sampled only from Hispanics. The model performance was evaluated using multiple parameters: accuracy, confusion matrix, receiver operating characteristics area under the curve (ROC AUC). For each model trained within the Random Forest ensemble, we obtained a vector representing the importance of each feature (i.e. gene). To identify the most relevant features (the relative importance - frequency - of each feature in a dataset when building a predictive model), we calculated the average contribution of each feature across all 1,000 models. Then, we ranked the features based on their average contribution, helping us pinpoint the most influential ones.

### Polytranscriptomics risk score

We generated a polygenic transcriptome risk score (PTRS) using several filters. Starting from all gene transcripts (58,954) and after removing pseudogenes and genes with zero expression level, we retained 48,356 genes. We used the ML findings to prioritize the gene list to be included in the PTRS: all the genes with a significant contribution (average weight >0, total *n=*43,718 for the agnostic model, and total *n=*43,635 for the aware model) were then employed in a logistic regression to estimate the weights (i.e. {3) within the training set, adjusting for the covariates (sex, age, RIN). The PTRS for each sample was constructed as the weighted sum of the individual’s Log_10_ of the normalized RNA-seq expression values of K identified genes (with beta different than zero):

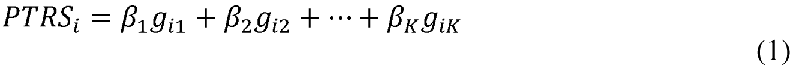

where β denotes weight and ***g*** denotes Log_10_ of the normalized RNA-seq gene expression values. Four different regression logistic models were implemented for AUC calculations:

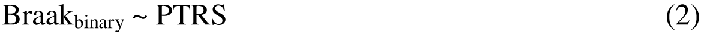

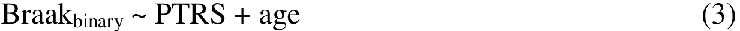

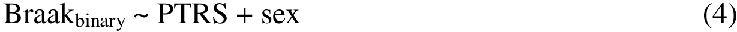

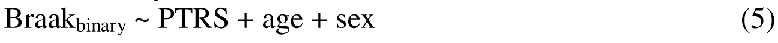

Finally, the prediction model with PTRS and covariates was implemented in the testing dataset and validated in ROS/MAP. The ROC-AUC was used to compare the models’ performances via the R package *pROC* (version 1.18.5). We used the R package *cutpoint* (version 1.1.2) to estimate the optimal cut points that maximizes the Youden-Index for determining the Braak binary outcome and validated performances using bootstrapping. Delong test was used to test significant difference between AUCs.

### Comparison of transcriptomics to human single nucleus dataset

To evaluate the RNA expressions of prioritized genes at the single nucleus/cell level, we used publicly available human single nucleus RNA sequencing (snRNAseq) dataset ^22^. Matrices were generated with 10X function of *Seurat* (version 4.1.3) R package ^23^. To generate a Seurat object, we filtered out any cells with less than 200 expressed genes, and with genes expressed in less than 3 cells. Following the normalization of the dataset, top 2,000 variable genes were used for further analyses. Anchors were identified by using *FindIntegrationAnchors*, and integration was performed using the *IntegrateData* functions of *DoubletFinde*r ^24^, while differential gene expressions were identified by *FindMarkers* function.

## Results

We included 565 brains in the Main model (mean age at death of 80.8±9.76 years old; 43.5% females; 96.1% with a primary neuropathological diagnosis of AD). The Columbia University New York Brain Bank contributed with the largest cohort, accounting for 46.2% of the total samples. 399 (70.6%) of the participants were self-reported as NHW, 113 (20.2%) as Hispanic, and 12 (2.1%) as African Americans. 85% of the participants received late-stage Braak neuropathological diagnosis (Braak > IV). **Table 1** reports the cohort demographics, while **Supplemental Fig. 1** shows the different sample sizes across Main model and model 2-4, depending on diagnosis selection and covariates availability. We included 1,095 samples in the validation dataset from ROS/MAP (mean age at death of 90.0±6.59 years old; 67.12% females; 98.5% NHW; 63.20% with a primary neuropathological diagnosis of AD; **Supplemental Table 1**). **Supplemental Fig.** 2 illustrates the first and second principal components of the gene expression counts of the individuals included in Main model, highlighted by AD status, continuous Braak, sex, and ethnicity.

**Table 1.** Demographics for MU-BRAIN.

|  | Descriptive statistics |
| --- | --- |
| Sample size | 565* |
| AGE (SD) | 80.82 (9.76) |
| SEX | 43.54% |
| Alzheimer's Disease – primary diagnosis | 96.11% |
| Mean PMI (SD) | 3.47 (15.66) |
| Mean RIN (SD) | 5.06 (1.16) |
| **Cohorts** |  |
| Columbia University | 46.20% |
| Mayo Clinic, Jacksonville | 12.21% |
| Indiana University (NCRAD) | 25.13% |
| University of Miami | 9.38% |
| University of California, San Diego | 6.02% |
| University of California, Davis | 1.06% |
| **Ethnicity\*** |  |
| Non-Hispanic Whites | 70.62% |
| Hispanics | 20.00% |
| African Americans | 2.12% |
| **Braak stage** |  |
| 0 | 0.53% |
| I | 1.06% |
| II | 1.06% |
| III | 3.19% |
| IV | 8.85% |
| V | 24.96% |
| VI | 60.35% |

\* 565 brains were used to build the Main model; 41 have missing ethnicity.

### Differential gene expression analysis

A total number of 1,630 and 480 genes were found significantly differentially expressed across the four models using the continuous or binary Braak phenotype, respectively. **Figure 1A** illustrates the differentially expressed genes identified by the Main model. Two four-way Venn diagrams (**Figure 2**) and correlation plots (**Supplemental Fig. 3**) confirmed the strong correlation across Main model and Models 2-4, either using binary or continuous Braak staging. **Table 2** and **Supplemental Table 2** include top DEG identified in the Main model (continuous and binary Braak, respectively). Top DEG include AD-known genes *BDNF* and *VGF* (*P_adj_* =1.41X 10^-4^ and *P_adj_*=3.44X 10^-7^ , respectively). An UpSet and a heatmap plots are also reported in **Supplemental Fig. 4**-**5**.

**Figure 1.**
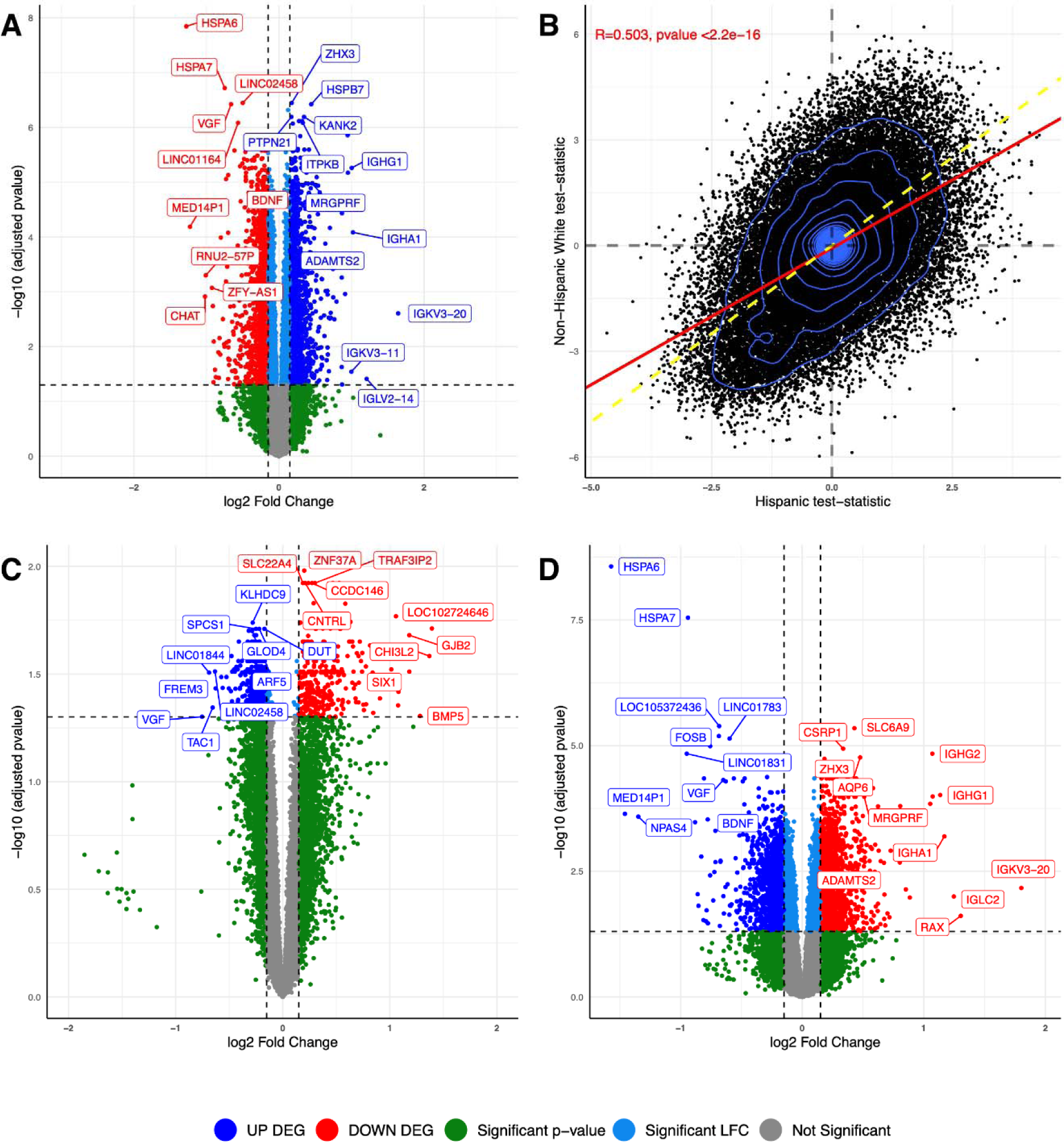
**(A)** Volcano plot of differentially expressed genes (Main model). **(B)** Correlation between NHW and Hispanic transcriptomes; points represent t-statistics from the ethnic-stratified differential expression analyses. Ethnic-stratified Volcano plots (**C**=Hispanics; **D**=NHW; Main model).

**Figure 2.**
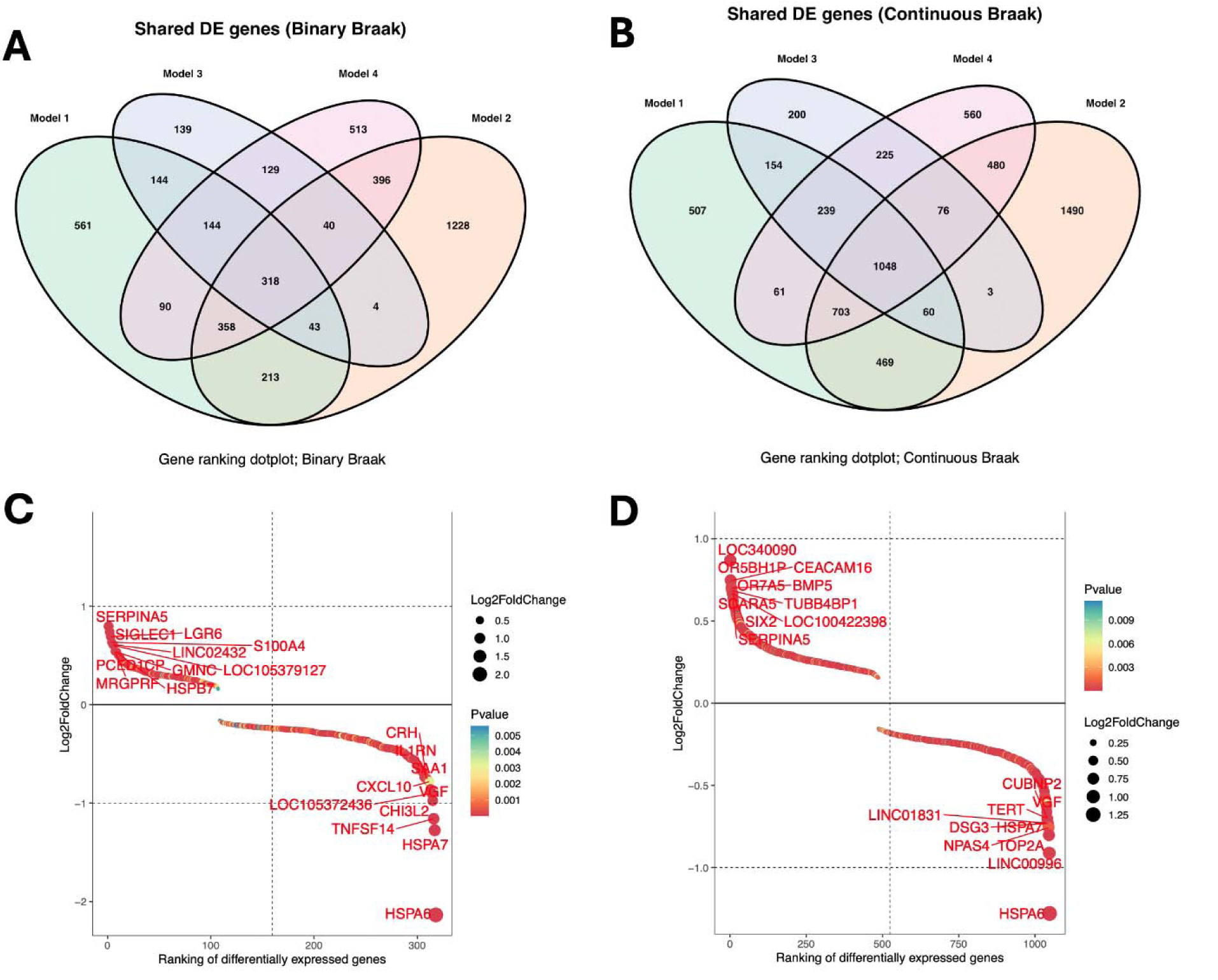
**(A) (B)** Venn diagrams of differentially expressed genes across Model 1-4 with binary and continuous Braak (Model 1=Main model). **(C) (D)** Gene ranking plots of the differentially expressed genes uniformly identified by Model 1-4 with binary and continuous Braak.

**Table 2.** Top DEG (p_adj_ < 1 x 10^-6^) associated with both continuous and binary Braak (Main model).

| **gene** | **Mean Exp** | **LFC** | **LFC SE** | **p** | **p<sub>adj</sub>** |
| --- | --- | --- | --- | --- | --- |
| **HSPA6** | 625.629 | -1.278 | 0.176 | 3.25e-13 | 1.43e-08 |
| **HSPA7** | 160.241 | -0.748 | 0.11 | 8.67e-12 | 1.91e-07 |
| **LINC02458** | 67.61 | -0.5 | 0.076 | 4.86e-11 | 3.57e-07 |
| **ZHX3** | 11671.853 | 0.176 | 0.027 | 3.69e-11 | 3.57e-07 |
| **HSPB7** | 133 | 0.445 | 0.068 | 6.17e-11 | 3.78e-07 |
| **VGF** | 1730.121 | -0.661 | 0.101 | 6.86e-11 | 3.78e-07 |
| **PTPN21** | 2039.003 | 0.171 | 0.027 | 2.04e-10 | 6.42e-07 |
| **KANK2** | 2279.701 | 0.345 | 0.054 | 1.86e-10 | 6.42e-07 |
| **ITPKB** | 28404.246 | 0.316 | 0.05 | 2.66e-10 | 7.65e-07 |
| **MTSS2** | 11148.713 | 0.276 | 0.044 | 2.78e-10 | 7.65e-07 |
| **TEKT2** | 101.495 | 0.328 | 0.052 | 3.36e-10 | 8.24e-07 |
| **LINC01164** | 104.189 | -0.564 | 0.09 | 3.34e-10 | 8.24e-07 |
| **TTC23** | 974.46 | 0.19 | 0.03 | 3.68e-10 | 8.54e-07 |

LFC= log<sub>2</sub> Fold Change; SE= standard error; p=p-value

To refine our findings, we prioritized 30 DEG that were replicated in ROS/MAP (**Table 3**), including: *ADAMTS2* (LFC=0.253, *P_adj_* =1.21X 10^-4^)*, VGF* (LFC=-0.661, *P_adj_* =3.78X 10^-7^), and *HSPB7* (LFC=0.445; *Padj*=3.78X 10^-7^ ). **Supplemental Table 3** presents the top DEG identified by the transethnic metanalysis, including *ADAMTS2* (LFC_meta_=0.265, *P_adj_*_-meta_=1.00X 10^-16^ )*, VGF* (LFC_meta_ =-0.332, *P_adj_*_-meta_=2.24X 10^-7^), and *HSPA6* (LFC_meta_ =-0.566; *P_adj_*_-meta_=1.17X 10^-O7^).

**Table 3.** Top MU-BRAIN DEG (p_adj_ < 1 x 10^-3^; Main model + RNA metrics) also replicated in ROS/MAP.

| Gene | MU-BRAIN |  |  |  | ROS/MAP |  |
| --- | --- | --- | --- | --- | --- | --- |
| | Mean Exp | LFC | LFC SE | $p_{\text{adj}}$ | LFC | $p_{\text{adj}}$ |
| *HSPB7* | 133 | 0.445 | 0.068 | 3.78e-07 | 0.164 | 0.00723 |
| *VGF* | 1730.121 | -0.661 | 0.101 | 3.78e-07 | -0.328 | 0.00156 |
| *ITPKB* | 28404.246 | 0.316 | 0.05 | 7.65e-07 | 0.155 | 0.000554 |
| *LINC01164* | 104.189 | -0.564 | 0.09 | 8.24e-07 | -0.223 | 0.00658 |
| *SLC6A9* | 1418.577 | 0.35 | 0.058 | 2.55e-06 | 0.172 | 7.08e-05 |
| *AQP6* | 102.784 | 0.413 | 0.069 | 2.55e-06 | 0.238 | 4.58e-05 |
| *SAMD4A* | 8062.731 | 0.295 | 0.05 | 3.59e-06 | 0.156 | 0.000238 |
| *GAREM2* | 2958.11 | 0.34 | 0.059 | 5.46e-06 | 0.248 | 6.6e-06 |
| *ANGPT1* | 402.304 | 0.395 | 0.069 | 5.46e-06 | 0.24 | 5.18e-05 |
| *TEAD2* | 197.351 | 0.422 | 0.074 | 5.46e-06 | 0.158 | 0.00526 |
| *CHST6* | 2096.031 | 0.379 | 0.067 | 6.71e-06 | 0.176 | 0.00286 |
| *MRGPRF* | 117.063 | 0.454 | 0.082 | 1.05e-05 | 0.263 | 7.01e-06 |
| *DDIT4L* | 588.868 | 0.402 | 0.073 | 1.06e-05 | 0.163 | 0.0198 |
| *IGFBP5* | 8972.946 | 0.438 | 0.081 | 1.29e-05 | 0.198 | 4.48e-05 |
| *LINC02552* | 253.266 | -0.419 | 0.078 | 1.56e-05 | -0.168 | 0.0155 |
| *NRIP2* | 661.458 | 0.185 | 0.034 | 1.58e-05 | 0.197 | 6.6e-06 |
| *NPNT* | 500.897 | 0.414 | 0.077 | 1.64e-05 | 0.31 | 5e-09 |
| *TRIP10* | 633.267 | 0.335 | 0.064 | 2.66e-05 | 0.181 | 0.000544 |
| *LINC00310* | 163.586 | 0.233 | 0.045 | 3.35e-05 | 0.255 | 0.0017 |
| *NAT16* | 41.22 | -0.423 | 0.082 | 3.36e-05 | -0.277 | 0.000144 |
| *A2ML1* | 999.685 | 0.275 | 0.055 | 4.99e-05 | 0.238 | 9.4e-05 |
| *CCDC102A* | 136.912 | 0.308 | 0.062 | 5.31e-05 | 0.176 | 0.00355 |
| *IQGAP3* | 108.579 | -0.329 | 0.067 | 6.95e-05 | -0.262 | 6.6e-06 |
| *ADAMTS2* | 1220.639 | 0.253 | 0.054 | 0.000121 | 0.329 | 1.06e-08 |
| *SLC4A11* | 241.876 | 0.26 | 0.055 | 0.000121 | 0.276 | 6.6e-06 |
| *APLN* | 1208.704 | 0.353 | 0.075 | 0.000122 | 0.215 | 0.000742 |
| *SMTN* | 1714.79 | 0.334 | 0.074 | 0.000207 | 0.17 | 0.00312 |
| *SLC38A2* | 12680.317 | 0.255 | 0.057 | 0.000273 | 0.234 | 1.96e-06 |
| *GIPR* | 187.961 | 0.442 | 0.1 | 0.000298 | 0.289 | 0.00693 |
| *GPCPD1* | 2137.119 | -0.226 | 0.053 | 0.000446 | -0.165 | 0.00076 |
| *EMP3* | 269.718 | 0.26 | 0.061 | 0.000448 | 0.172 | 0.000751 |
| *LINC02217* | 774.81 | -0.224 | 0.053 | 0.00053 | -0.155 | 0.00497 |
| *TNFRSF18* | 26.298 | -0.345 | 0.082 | 0.000586 | -0.171 | 0.0354 |
| *PRELP* | 2316.061 | 0.24 | 0.057 | 0.000606 | 0.227 | 1.87e-06 |
| *SLC7A2* | 4964.733 | 0.295 | 0.072 | 0.000787 | 0.151 | 0.0254 |

LFC= $\log_2$ Fold Change; SE= standard error; p=p-value

### Ethnic-stratified analyses

Overall, we observed high concordance between Hispanic and NHW transcriptome (Spearman’s rho=0.503; p < 2.2 X 10^-16^; **Figure 1B**). **Table 4** displays the top DEG identified in the ethnic-stratified models, while **Supplemental Table 4** outlines top DEG consistently identified in both NHW and Hispanics, i.e. genes that pinpoint to universal loci underlining AD pathology regardless of ethnicity **(Figures 1C-D**). *VGF* was identified in both Hispanics and NHW groups (Hispanic: LFC=-0.753, *P_adj_* =0.0499; NHW: LFC=-0.650, *P_adj_* =4.84X 10^-5^ ). While several *HSP* family genes were consistently differentially expressed within Hispanics and NHW (e.g. *HSPB7*; Hispanic: LFC=0.629, *P_adj_* =0.0224; NHW: LFC=0.408, *P_adj_* =1.80X 10^-4^), others were uniquely differentially expressed within NHW (e.g. *HSPA6*; NHW: LFC=-1.57, *P_adj_* =2.72X 10^-9^ ; Hispanic: LFC=0.467, *P_adj_*=0.449). *CHI3L2* was found uniquely overexpressed in Hispanics (LFC=1.370; *P_adj_* =0.026) but not in NHW (LFC=-0.47, *P_adj_* =0.105). The ethnicity interaction model prioritized two DEG: *TNFSF14* (LFC=-0.506; *P_adj_* =0.00268) and *SPOCD1* (LFC=-0.33; *P_adj_* =0.00319).

**Table 4.** Top DEG (p_adj;Hispanics_ < 1.5 x 10^-2^; p_adj;NHW_ < 1.5 x 10^-3^) stratified by ethnicity (Main model).

| gene | Mean Exp | LFC | LFC SE | p | $p_{\text{adj}}$ |
| --- | --- | --- | --- | --- | --- |
| **Hispanics** |  |  |  |  |  |
| **ZNF37A** | 4175.035 | 0.201 | 0.041 | 7.33e-07 | 0.0105 |
| **SLC22A4** | 446.889 | 0.189 | 0.04 | 2.24e-06 | 0.0119 |
| **TRAF3IP2** | 2634.819 | 0.301 | 0.063 | 2e-06 | 0.0119 |
| **CCDC146** | 850.704 | 0.274 | 0.059 | 3.42e-06 | 0.0119 |
| **CNTRL** | 3564.157 | 0.21 | 0.046 | 5.9e-06 | 0.0119 |
| **CDKN2B** | 256.352 | 0.468 | 0.103 | 5.97e-06 | 0.0119 |
| **RESF1** | 4267.864 | 0.24 | 0.053 | 5.23e-06 | 0.0119 |
| **CNTLN** | 1027.817 | 0.289 | 0.065 | 8.08e-06 | 0.0148 |
| **PLP2** | 110.09 | 0.584 | 0.131 | 8.79e-06 | 0.0149 |
| **NHW** |  |  |  |  |  |
| **UQCRC2** | 4604.066 | -0.132 | 0.028 | 1.9e-06 | 0.000359 |
| **MED10** | 739.921 | -0.129 | 0.028 | 3.1e-06 | 0.000454 |
| **SPCS1** | 889.946 | -0.139 | 0.03 | 4.27e-06 | 0.00054 |
| **CIAPIN1** | 958.817 | -0.124 | 0.028 | 1.08e-05 | 0.000866 |
| **OCIAD1** | 4124.498 | -0.137 | 0.032 | 1.39e-05 | 0.000978 |
| **RAMACL** | 481.161 | 0.121 | 0.028 | 1.64e-05 | 0.00106 |
| **VDAC2** | 2107.145 | -0.135 | 0.032 | 2.27e-05 | 0.00129 |
| **NDUFB8** | 641.275 | -0.117 | 0.028 | 2.38e-05 | 0.00132 |

LFC= $\log_2$ Fold Change; SE= standard error; p=p-value; NHW= non-Hispanic Whites

### Gene set variation analysis (GSVA) and functional enrichment analysis

GSVA (Main model) identified a 118 differentially enriched gene sets (**Supplemental Fig. 6A)**. A heatmap plot of GSVA enrichment scores associated with the top differentially expressed gene set is presented in **Supplemental Fig. 7**. **Supplemental Table 5** shows the top differentially enriched gene sets identified by GSVA in association with the Main model.

Figure 3A illustrates the most significantly overrepresented gene lists with GSEA using WikiPathways, where the pathway “*Alzheimer disease*” (WP5124; *Padj*=4.24 X 10^-6^ ) and “*Alzheimer disease and miRNA effects*” (WP2059; *P_adj_* =4.24X 10^-6^) have the largest counts of genes among the significant gene lists. Gene-concept networks (**Supplemental** Fig. 8) were also presented to demonstrate the relationships between genes and the overarching biological concepts/pathways. Several genes exhibited strong ties to concept “*TYROBP* causal network in microglia” (WP3945; *P_adj_* =1.68X 10^-8^).

**Figure 3.**
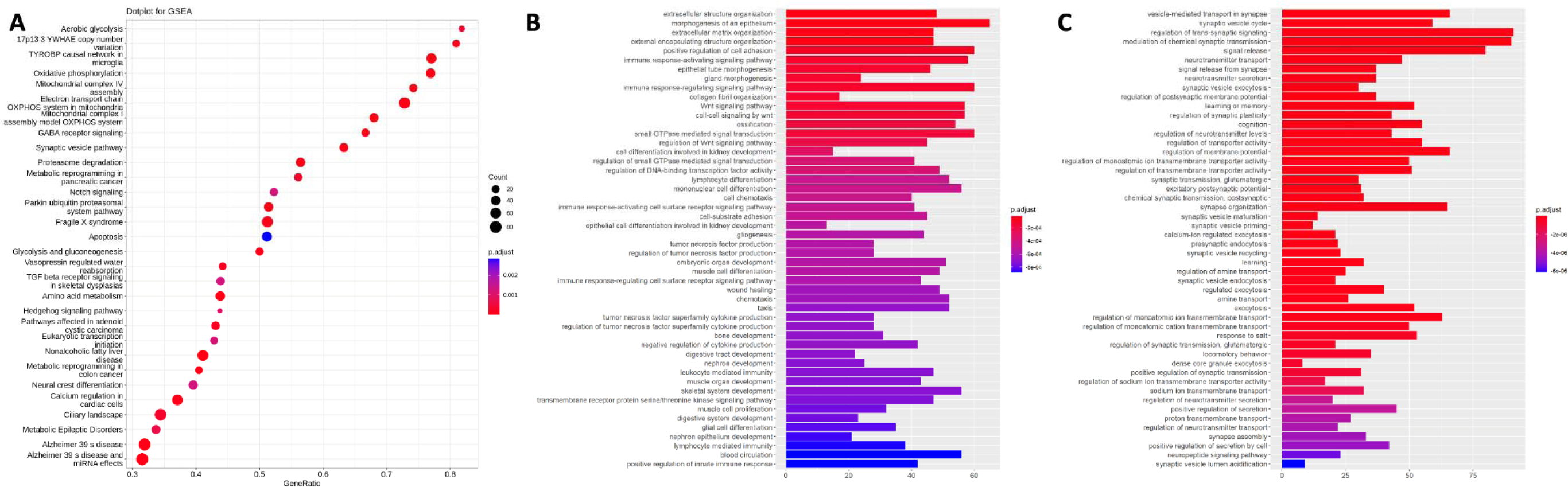
Results of GSVA and functional enrichment analysis. **(A)** Top GSEA Wikipathways from functional enrichment analysis; **(B)** Top GO Enriched Pathways for up-regulated genes; **(C)** Top GO Enriched Pathways for down-regulated genes.

GO enrichment analyses were implemented separately in up-regulated and down-regulated genes. Up-regulated genes were enriched in several pathways, including “*Immune response activation signal pathways”* and “*extracellular structure organization*” (*P_adj_* =1.16X 10^-4^ and *P_adj_* =2.27X 10^-5^, respectively; Figure 3B). Among the top 50 pathways of down-regulated genes, 21 were

found to be enriched in synapse-related processes (e.g. “*Vesicle-mediated transport in synapse*”, “*regulation of trans-synaptic signaling*”, “*modulation of chemical synaptic transmission*”; *P_adj_*< 1 X 10^-17^ ; Figure 3C). The remaining pathways included neurotransmitter, learning and cognitive pathways (“*neurotransmitter transport*”, “*learning or memory”*, and “*cognition*”; all *P_adj_* <1X 10^-8^; Figure 3C). **Supplemental** Figures 9 **A-B** reconstruct networks of genes and their corresponding biological pathways.

### Stratified analyses by ethnic group in GSVA and functional enrichment analysis

**Supplemental** Fig. 10A-**B** display the heatmaps of GSVA enrichment scores for the Main model stratified by Hispanics and NHW: 677 gene sets were differentially enriched within Hispanics, and 145 in NHW. **Supplemental** Fig. 11 and **Supplemental Table 6** present the differentially enriched gene sets within each ethnic subsample. The pathway “*Alzheimer disease*”, “*Alzheimer disease and miRNA effects*”, and “*TYROBP causal network in microglia*” were consistently identified as top significantly overrepresented gene lists in Main model for both Hispanics and NHW (**Supplemental** Fig. 12 **A&B**). The GO enrichment analysis in Main model stratified by Hispanic and NHW groups were provided in **Supplemental** Fig. 13&**14 A-D**. **Supplemental Tables 7**&**9** display the top pathways uniquely associated with either Hispanic or NHW cohorts, as identified through GSEA and GO enrichment analysis in Main model stratified by ethnicity. **Supplemental Tables 8**,**10** present the top pathways common to both ethnicities.

### Single-nucleus expression of prioritized genes

Our analyses identified several concordant DEG between bulk RNAseq and snRNAseq (**Supplemental** Fig. 15) despite their inherent technical differences ^25^, including *HSPA6*, found downregulated in microglia (LFC=-1.488, *P_adj_* =1.1E-07), *PCSK1* found downregulated in neurons (LFC=-0.034, *P_adj_* =4.2E-06) and endothelial cells (LFC=-1.050, *P_adj_* =3.9E-05), and *SLC7A2* found upregulated in astroglia (LFC=0.630, *P_adj_* =2.1E-06).

### Phenotype prediction using machine learning algorithm

In the “Ethnicity-Agnostic” Model, the random testing sample showed an AUC=0.74 (**Supplemental** Fig. 16A). In the “Ethnicity-Aware” Model, the Hispanics + AA testing sample showed an AUC=0.70 (**Supplemental** Fig. 16B). The genes identified as most relevant for prediction (“feature importance”; average weight >0; total of 43,718 for the Ethnicity-Agnostic model and 43,635 for the Ethnicity-Aware model) were then selected to compile a priority gene list for subsequent use in PTRS (**Supplemental Table 11)**.

### Polytranscriptomics risk score

After prioritizing genes using the random forest ML and logistic regression, the PTRS was constructed using 256 and 284 genes (Agnostic and Aware models, respectively; **Supplemental** Fig. 17). The PTRS effectively classified brains in low vs. high Braak stage in both the “Ethnicity-Agnostic” and “Ethnicity-Aware” model (ANOVA *P* =7.5X 10^-4^, and *P* =2.5 X 10^-3^, respectively). The Violin plots for each model are shown in Figures 4A-B.

**Figure 4.**
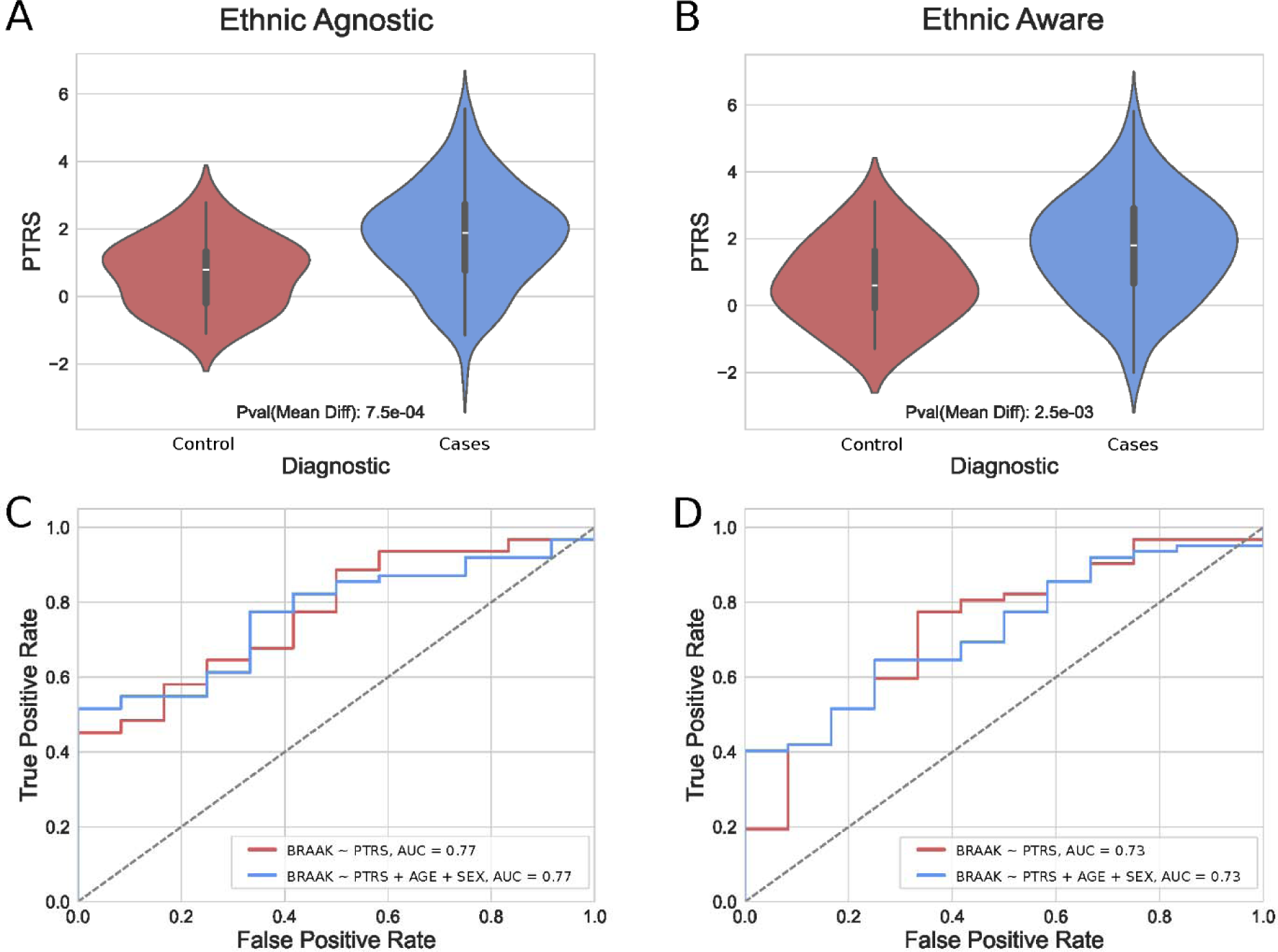
PTRS in high Braak (1) vs low Braak (0) stages within the testing sample for PTRS’s AUCs (plus covariates). **(A)** “Ethnicity-Agnostic” . **(B)** “Ethnicity-Aware” . **(C)** Ethnicity-Agnostic. **(D)** Ethnicity-Aware models.

The model showed an AUC=0.78 in the “Ethnicity-Agnostic” model, and AUC=0.73 in the “Ethnicity-Aware” model. We did not observe significant differences in terms of performance between the two models (*P*_DeLong_ = 0.4; Figures 4A-B). Additionally, the “Ethnicity-Agnostic” PTRS was validated in ROS/MAP (**Supplemental** Fig. 18), where it effectively classified brains into low vs. high Braak stage (ANOVA *P* =5.2X 10^-3^).

## Discussion

By utilizing a large number of brain samples as well as including individuals from various ethnic backgrounds, we implemented a range of conventional and innovative methods to explore the association between transcriptomic patterns and the manifestation of AD-related hallmarks, specifically Braak staging. Additionally, we examined how ethnicity might influence this association, shedding light on the potential role of ethnicity in modifying the impact of AD-related hallmarks.

Several genes were consistently differentially expressed across various models, such as *VGF* and members of the *HSP* family emerging as strongly associated with higher Braak stages. The consistent identification of *VGF,* widely reported in previous studies ^26–29^, underscores its potential significance as a key regulator in AD mechanisms. *VGF* is downregulated in neurons in AD brains and in amyloid toxicity model of zebrafish ^30,31^, indicating an evolutionarily conserved disease mechanism. The *HSP70* family is a class of highly abundant and widely expressed chaperone proteins, encompassing up to 17 genes and 30 pseudogenes. *HSP70*, which is known to regulate protein misfolding including tau levels and toxicity, colocalizes in AD brains with Aβ plaques and takes part in the neuroprotective response to suppress Aβ aggregation. A recent study revealed expression levels of *HSPA1A* and *HSPA2* were significantly increased in AD, while *HSPA8* significantly decreased^32^. *VGF* has also been identified as a key regulator of AD-associated network, where its overexpression is associated with partial rescue of Aβ- mediated memory impairment and neuropathology, suggesting its potential protective role against AD progression.^33^ The validation provided by ROS/MAP further enhances the robustness of our results. In the meta-analysis, genes identified as differentially expressed in both MU- BRAIN and ROS/MAP exhibited identical directions of disexpression, underscoring a consistent regulatory impact on Braak staging across ethnicities and studies.

*HSPA6* was specifically downregulated in NHW in bulk and single nucleus data (Figure 1A, **Table 3, Supplemental** Fig. 15). Previous studies in mice^34^, zebrafish^30,31^ and fruit fly ^35^ have confirmed the relevance of *Hsp70* genes to age-related neurodegeneration and amyloid toxicity, suggesting an evolutionarily conserved role of *HSP70* proteins in resilience against neurodegeneration. In human single nucleus data, we found that *HSPA6* is significantly downregulated in AD in microglial cells (**Supplemental** Fig. 15), supporting the critical role of immune system in clearing toxic protein aggregates in the brain. Consistent with this finding, we identified several GO-terms and pathways related to microglial activity. For instance, the “TYROBP causal network in microglia” pathway has been previous indicated to be pivotal in microglial activation in AD^36^. *TYROBP/DAP12* has indeed been identified as a key AD modulator, when its suppression leads to elevated tau pathology and cognitive deficits, suggesting its role in mediating tau toxicity through microglial and oligodendrocyte interactions^37^. Additionally, the GSEA analysis underscores the significance of the “*Alzheimer disease and miRNA effects*” pathway, echoing recent expert-curated data from a comprehensive study that enhanced understanding of key proteins and interactions pivotal to AD research^38^. The identification of the "*Alzheimer disease*" pathway aligns with an independent study that employed a network-based strategy to discern anti-AD compounds, particularly emphasizing the apoptosis of neuron cells^39^. Studies have further indicated that the loss of synaptic structure and function is a hallmark among individuals with AD^40,41^. In our study, we found pathways related to neuronal connectivity and function (**Supplemental** Fig. 7**,8**), and further supported by our findings of downregulation of *GPCPD1* (**Table 3, Supplemental** Fig. 15), a gene required for formation of cholinergic neurotransmitters that are reduced in AD ^42,43^ and reduced *RPH3A*, a critical regulator of neuronal synapse potentiation, neurotransmission and memory ^44^. Downregulated choline acetyltransferase gene *CHAT,* which is required for production of acetylcholine neurotransmitter ^45^, also supports these findings.

The pathway “*activation of immune responses*” identified in GO enrichment analysis has also been highlighted in previous studies^46^. Notably, two pathways related to extracellular matrix (ECM) organization, i.e. “*extracellular structure organization*” and “*extracellular matrix organization*”, have also been shown to be pivotal in organizing ECM in neurodegenerative diseases^47^. Consistently, the upregulated genes *KANK2* (**Table 3**) is required for cell polarization to regulate cell migration in response to extracellular matrix stiffness ^48^. Similarly, *ADAMTS2,* that we found upregulated in AD, and consistently reported in AD genome-wide association study (GWAS) studies, is a matrix metalloproteinase required for production of functional collagen residues in the extracellular matrix and potentially leading to extracellular stiffening that affects cellular signaling. Since ECM is critical for disease progression and pathophysiology of AD ^49,50^, ancestry-specific alterations in expression of ECM components can provide new drug targets. Interestingly, *ADAMTS2* was ranked as the #1 gene in our random forest algorithm.

The inclusion of brains from different ethnic groups, with a specific representation of individuals of Hispanic/Latino ethnicity, identified ethnic-specific transcriptome profiles. Top differentially expressed gene in Hispanic, *TNFSF14*, has been previously identified as a protective factor for vascular dementia in women^51^. *SPOCD1* has also been linked to DNA methylation, which has been shown to be altered in AD^52,53^. Lastly, *CHI3L2* has been suggested as potential biomarker in glioma^54^, a condition which frequency varies between Hispanics and NHW ^55,56^.

Multiple studies have shown significant differences in gene expression profiles across populations, with particular emphasis in immune response and metabolism-related gene pools^57^. These differences, which are influenced by genomic profiles and are often tissue-specific, highlight the complexity of population-specific gene regulation. Despite these findings, it is important to note our data might not be generalizable to the general population. Indeed, autopsy-based collections have numerous selection biases. It is also imperative to denote race and ethnicity are social constructs and can have numerous intertwined variables of which may relate to differences found^58^. Nevertheless, when NHW were employed to train the machine learning algorithm and Hispanics for testing (Ethnicity-Aware model), we found the model to perform comparably to the Ethnicity-agnostic one. This is in line with our recently published investigation where we showed a strong correlation between Hispanic and NHW transcriptome profiles^59^, ultimately confirming that the brain region used to extract RNA was more relevant than the ethnic group the brain belonged to. Additional proofs of the optimal transferability across ethnicities comes from a recent publication showing that gene expression signatures of aging in peripheral blood are shared between East Asian and NHW populations, with specific hub genes like *NUDT7* and *OXNAD1*, decreasing in expression with age^60^. This finding reinforces the universality of aging biomarkers across ethnicities, which further validates our study.

Polygenic risk scores (PRS) play a crucial role in bridging the gap between GWAS and their application in clinical settings. Recently, new approaches have been developed to generate risk scores employing OMICS data, although training samples are often limited to a small subset of genes^61^. To this end, we introduced a polytranscriptomic risk score that was trained on the entire spectrum of genes to predict AD-related pathology, marking a significant advancement in our ability to predict and stratify samples for further investigations. Indeed, utilizing a comprehensive set of genes has been demonstrated to be more effective in enhancing predictions in AD studies^62^. Our results ultimately suggest that the PTRS matches or even outperforms the predictive accuracy of traditional PRS^63^. Notably, our score showed comparable performances between ’Ethnicity-Agnostic’ and ’Ethnicity-Aware’ models, contrary to previous investigations reporting significant differences across ethnicities^64^. The outcomes of our study highlight a potential implementation of PTRS in developing predictive models for AD pathology independently of ethnic backgrounds. This assertion was reinforced by deploying our PTRS in the ROS/MAP cohort, revealing its ability to distinguish between low and high Braak stages.

The current study has limitation, notably the underrepresentation of African American and Asian samples. Furthermore, there is no consideration or comparison work on the diversity within Hispanic samples (e.g. Caribbean Hispanics, Mexicans, and others)^65^. Moreover, while our research has made considerable strides in characterizing the transcriptome associated with Braak staging, the resolution of insights is inherently limited using bulk RNA sequencing data. Despite applying bioinformatic tools to mitigate the batch effects, variation in procedures among AD centers may cause potential bias in the current study. Future efforts will focus on single-cell data of diverse ethnicities to enable a more refined analysis at the cellular level, thereby enhancing the significance of our findings.

In conclusion, the expansion brought by MU-BRAIN to include a broader representation of ethnic groups and the advancement of our methodological approach to a single-cell resolution would undoubtedly augment the depth and applicability of our research in uncovering the molecular underpinnings of AD diverse populations.

## Data availability

Please contact the correspondence author for data related to this study.

## Supplementary material

Supplementary material is available at *Brain* online.

## REFERENCES

1. Braak H, Braak E. Neuropathological stageing of Alzheimer-related changes. Acta Neuropathol. 1991;82(4):239–59. doi:10.1007/BF00308809

2. Bagyinszky E, Giau VV, An SA. Transcriptomics in Alzheimer’s Disease: Aspects and Challenges. Int J Mol Sci. May 15 2020;21(10)doi:10.3390/ijms21103517

3. Annese A, Manzari C, Lionetti C, et al. Whole transcriptome profiling of Late-Onset Alzheimer’s Disease patients provides insights into the molecular changes involved in the disease. Sci Rep. Mar 9 2018;8(1):4282. doi:10.1038/s41598-018-22701-2

4. Rodriguez-Esteban R, Jiang X. Differential gene expression in disease: a comparison between high-throughput studies and the literature. BMC Med Genomics. Oct 11 2017;10(1):59. doi:10.1186/s12920-017-0293-y

5. Reitz C, Mayeux R. Genetics of Alzheimer’s disease in Caribbean Hispanic and African American populations. Biol Psychiatry. Apr 1 2014;75(7):534–41. doi:10.1016/j.biopsych.2013.06.003

6. Chin AL, Negash S, Hamilton R. Diversity and disparity in dementia: the impact of ethnoracial differences in Alzheimer disease. Alzheimer Dis Assoc Disord. Jul-Sep 2011;25(3):187–95. doi:10.1097/WAD.0b013e318211c6c9

7. Hill CV, Perez-Stable EJ, Anderson NA, Bernard MA. The National Institute on Aging Health Disparities Research Framework. Ethn Dis. Aug 7 2015;25(3):245–54. doi:10.18865/ed.25.3.245

8. Balls-Berry JJE, Babulal GM. Health Disparities in Dementia. Continuum (Minneap Minn). Jun 1 2022;28(3):872–884. doi:10.1212/CON.0000000000001088

9. Therriault J, Pascoal TA, Lussier FZ, et al. Biomarker modeling of Alzheimer’s disease using PET-based Braak staging. Nat Aging. Jun 2022;2(6):526–535. doi:10.1038/s43587-022-00204-0

10. Macedo AC, Tissot C, Therriault J, et al. The Use of Tau PET to Stage Alzheimer Disease According to the Braak Staging Framework. J Nucl Med. Aug 2023;64(8):1171–1178. doi:10.2967/jnumed.122.265200

11. Montine TJ, Phelps CH, Beach TG, et al. National Institute on Aging-Alzheimer’s Association guidelines for the neuropathologic assessment of Alzheimer’s disease: a practical approach. Acta Neuropathol. Jan 2012;123(1):1–11. doi:10.1007/s00401-011-0910-3

12. Zhang Y, Parmigiani G, Johnson WE. ComBat-seq: batch effect adjustment for RNA-seq count data. NAR Genom Bioinform. Sep 2020;2(3):lqaa078. doi:10.1093/nargab/lqaa078

13. Liu Q, Shepherd B, Li C. PResiduals: an R package for residual analysis using probability-scale residuals. Journal of Statistical Software. 2020;94:1–27.

14. Love MI, Huber W, Anders S. Moderated estimation of fold change and dispersion for RNA-seq data with DESeq2. Genome Biol. 2014;15(12):550. doi:10.1186/s13059-014-0550-8

15. Lee AJ, Ma Y, Yu L, et al. Multi-region brain transcriptomes uncover two subtypes of aging individuals with differences in Alzheimer risk and the impact of APOEepsilon4. bioRxiv. Jan 25 2023; doi:10.1101/2023.01.25.524961

16. Van der Auwera GA, O’Connor BD, Safari aORMC. Genomics in the cloud : using Docker, GATK, and WDL in Terra. First edition. ed. O’Reilly Media; 2020.

17. Willer CJ, Li Y, Abecasis GR. METAL: fast and efficient meta-analysis of genomewide association scans. Bioinformatics. Sep 1 2010;26(17):2190–1. doi:10.1093/bioinformatics/btq340

18. Hanzelmann S, Castelo R, Guinney J. GSVA: gene set variation analysis for microarray and RNA-seq data. BMC Bioinformatics. Jan 16 2013;14:7. doi:10.1186/1471-2105-14-7

19. Liberzon A, Subramanian A, Pinchback R, Thorvaldsdottir H, Tamayo P, Mesirov JP. Molecular signatures database (MSigDB) 3.0. Bioinformatics. Jun 15 2011;27(12):1739–40. doi:10.1093/bioinformatics/btr260

20. Ritchie ME, Phipson B, Wu D, et al. limma powers differential expression analyses for RNA-sequencing and microarray studies. Nucleic Acids Res. Apr 20 2015;43(7):e47. doi:10.1093/nar/gkv007

21. Yu G, Wang LG, Han Y, He QY. clusterProfiler: an R package for comparing biological themes among gene clusters. OMICS. May 2012;16(5):284–7. doi:10.1089/omi.2011.0118

22. Lau SF, Cao H, Fu AKY, Ip NY. Single-nucleus transcriptome analysis reveals dysregulation of angiogenic endothelial cells and neuroprotective glia in Alzheimer’s disease. Proc Natl Acad Sci U S A. Oct 13 2020;117(41):25800–25809. doi:10.1073/pnas.2008762117

23. Stuart T, Butler A, Hoffman P, et al. Comprehensive Integration of Single-Cell Data. Cell. Jun 13 2019;177(7):1888–1902 e21. doi:10.1016/j.cell.2019.05.031

24. McGinnis CS, Murrow LM, Gartner ZJ. DoubletFinder: Doublet Detection in Single-Cell RNA Sequencing Data Using Artificial Nearest Neighbors. Cell Syst. Apr 24 2019;8(4):329–337 e4. doi:10.1016/j.cels.2019.03.003

25. Ziegenhain C, Vieth B, Parekh S, et al. Comparative Analysis of Single-Cell RNA Sequencing Methods. Mol Cell. Feb 16 2017;65(4):631–643 e4. doi:10.1016/j.molcel.2017.01.023

26. Hendrickson RC, Lee AY, Song Q, et al. High Resolution Discovery Proteomics Reveals Candidate Disease Progression Markers of Alzheimer’s Disease in Human Cerebrospinal Fluid. PLoS One. 2015;10(8):e0135365. doi:10.1371/journal.pone.0135365

27. Carrette O, Demalte I, Scherl A, et al. A panel of cerebrospinal fluid potential biomarkers for the diagnosis of Alzheimer’s disease. Proteomics. Aug 2003;3(8):1486–94. doi:10.1002/pmic.200300470

28. Brinkmalm G, Sjodin S, Simonsen AH, et al. A Parallel Reaction Monitoring Mass Spectrometric Method for Analysis of Potential CSF Biomarkers for Alzheimer’s Disease. Proteomics Clin Appl. Jan 2018;12(1)doi:10.1002/prca.201700131

29. El Gaamouch F, Audrain M, Lin WJ, et al. VGF-derived peptide TLQP-21 modulates microglial function through C3aR1 signaling pathways and reduces neuropathology in 5xFAD mice. Mol Neurodegener. Jan 10 2020;15(1):4. doi:10.1186/s13024-020-0357-x

30. Cosacak MI, Bhattarai P, De Jager PL, Menon V, Tosto G, Kizil C. Single Cell/Nucleus Transcriptomics Comparison in Zebrafish and Humans Reveals Common and Distinct Molecular Responses to Alzheimer’s Disease. Cells. 2022;11:1807. 10.3390/cells11111807

31. Cosacak MI, Bhattarai P, Reinhardt S, et al. Single-Cell Transcriptomics Analyses of Neural Stem Cell Heterogeneity and Contextual Plasticity in a Zebrafish Brain Model of Amyloid Toxicity. Cell Rep. Apr 23 2019;27(4):1307–1318 e3. doi:10.1016/j.celrep.2019.03.090

32. Dong Y, Li T, Ma Z, Zhou C, Wang X, Li J. HSPA1A, HSPA2, and HSPA8 Are Potential Molecular Biomarkers for Prognosis among HSP70 Family in Alzheimer’s Disease. Dis Markers. 2022;2022:9480398. doi:10.1155/2022/9480398

33. Beckmann ND, Lin WJ, Wang M, et al. Multiscale causal networks identify VGF as a key regulator of Alzheimer’s disease. Nat Commun. Aug 7 2020;11(1):3942. doi:10.1038/s41467-020-17405-z

34. Lackie RE, Razzaq AR, Farhan SMK, et al. Modulation of hippocampal neuronal resilience during aging by the Hsp70/Hsp90 co-chaperone STI1. J Neurochem. Jun 2020;153(6):727–758. doi:10.1111/jnc.14882

35. Martin-Pena A, Rincon-Limas DE, Fernandez-Funez P. Engineered Hsp70 chaperones prevent Abeta42-induced memory impairments in a Drosophila model of Alzheimer’s disease. Sci Rep. Jul 2 2018;8(1):9915. doi:10.1038/s41598-018-28341-w

36. Haure-Mirande JV, Audrain M, Ehrlich ME, Gandy S. Microglial TYROBP/DAP12 in Alzheimer’s disease: Transduction of physiological and pathological signals across TREM2. Mol Neurodegener. Aug 24 2022;17(1):55. doi:10.1186/s13024-022-00552-w

37. Luo W, Che H, Fan L, et al. DAP12 deficiency alters microglia-oligodendrocyte communication and enhances resilience against tau toxicity. bioRxiv. 2023:2023.10.26.563970. doi:10.1101/2023.10.26.563970

38. Breuza L, Arighi CN, Argoud-Puy G, et al. A Coordinated Approach by Public Domain Bioinformatics Resources to Aid the Fight Against Alzheimer’s Disease Through Expert Curation of Key Protein Targets. J Alzheimers Dis. 2020;77(1):257–273. doi:10.3233/JAD-200206

39. Li B, Wu YR, Li L, Liu Y, Yan ZY. A Novel Based-Network Strategy to Identify Phytochemicals from Radix Salviae Miltiorrhizae (Danshen) for Treating Alzheimer’s Disease. Molecules. Jul 12 2022;27(14)doi:10.3390/molecules27144463

40. Spires-Jones TL, Hyman BT. The intersection of amyloid beta and tau at synapses in Alzheimer’s disease. Neuron. May 21 2014;82(4):756–71. doi:10.1016/j.neuron.2014.05.004

41. John A, Reddy PH. Synaptic basis of Alzheimer’s disease: Focus on synaptic amyloid beta, P-tau and mitochondria. Ageing Res Rev. Jan 2021;65:101208. doi:10.1016/j.arr.2020.101208

42. Bakken TE, Roddey JC, Djurovic S, et al. Association of common genetic variants in GPCPD1 with scaling of visual cortical surface area in humans. Proc Natl Acad Sci U S A. Mar 6 2012;109(10):3985–90. doi:10.1073/pnas.1105829109

43. Ferreira-Vieira TH, Guimaraes IM, Silva FR, Ribeiro FM. Alzheimer’s disease: Targeting the Cholinergic System. Curr Neuropharmacol. 2016;14(1):101–15. doi:10.2174/1570159x13666150716165726

44. Deak F, Shin OH, Tang J, et al. Rabphilin regulates SNARE-dependent re-priming of synaptic vesicles for fusion. EMBO J. Jun 21 2006;25(12):2856–66. doi:10.1038/sj.emboj.7601165

45. Scacchi R, Gambina G, Moretto G, Corbo RM. Variability of AChE, BChE, and ChAT genes in the late-onset form of Alzheimer’s disease and relationships with response to treatment with Donepezil and Rivastigmine. Am J Med Genet B Neuropsychiatr Genet. Jun 5 2009;150B(4):502-7. doi:10.1002/ajmg.b.30846

46. Wang MM, Miao D, Cao XP, Tan L, Tan L. Innate immune activation in Alzheimer’s disease. Ann Transl Med. May 2018;6(10):177. doi:10.21037/atm.2018.04.20

47. Soleman S, Filippov MA, Dityatev A, Fawcett JW. Targeting the neural extracellular matrix in neurological disorders. Neuroscience. Dec 3 2013;253:194–213. doi:10.1016/j.neuroscience.2013.08.050

48. Wang T, Hamilla S, Cam M, Aranda-Espinoza H, Mili S. Extracellular matrix stiffness and cell contractility control RNA localization to promote cell migration. Nat Commun. Oct 12 2017;8(1):896. doi:10.1038/s41467-017-00884-y

49. Yang AC, Vest RT, Kern F, et al. A human brain vascular atlas reveals diverse mediators of Alzheimer’s risk. Nature. Mar 2022;603(7903):885-892. doi:10.1038/s41586-021-04369-3

50. Sun Y, Xu S, Jiang M, et al. Role of the Extracellular Matrix in Alzheimer’s Disease. Front Aging Neurosci. 2021;13:707466. doi:10.3389/fnagi.2021.707466

51. Lee C, Kim Y. Complex genetic susceptibility to vascular dementia and an evidence for its underlying genetic factors associated with memory and associative learning. Gene. Mar 2013;516(1):152–157. doi:10.1016/j.gene.2012.12.032

52. Yokoyama AS, Rutledge JC, Medici V. DNA methylation alterations in Alzheimer’s disease. Environ Epigenet. May 2017;3(2):dvx008. doi:10.1093/eep/dvx008

53. Zoch A, Auchynnikava T, Berrens RV, et al. SPOCD1 is an essential executor of piRNA-directed de novo DNA methylation. Nature. Aug 2020;584(7822):635-639. doi:10.1038/s41586-020-2557-5

54. Qian W, Wang Q, Zhang C, Zhu J, Zhang Q, Luo C. M2 macrophage marker CHI3L2 could serve as a potential prognostic and immunological biomarker in glioma by integrated single-cell and bulk RNA-Seq analysis. J Gene Med. Sep 2023;25(9):e3523. doi:10.1002/jgm.3523

55. Lehrer S. Glioma and Alzheimer’s Disease. J Alzheimers Dis Rep. Dec 14 2018;2(1):213–218. doi:10.3233/ADR-180084

56. Walsh KM, Neff C, Bondy ML, et al. Influence of county-level geographic/ancestral origin on glioma incidence and outcomes in US Hispanics. Neuro Oncol. Feb 14 2023;25(2):398–406. doi:10.1093/neuonc/noac175

57. Daca-Roszak P, Zietkiewicz E. Transcriptome variation in human populations and its potential application in forensics. J Appl Genet. Nov 2019;60(3-4):319–328. doi:10.1007/s13353-019-00510-1

58. Nguyen ML, Huie EZ, Whitmer RA, George KM, Dugger BN. Neuropathology Studies of Dementia in US Persons other than Non-Hispanic Whites. Free Neuropathol. 2022;3doi:10.17879/freeneuropathology-2022-3795

59. Felsky D, Santa-Maria I, Cosacak MI, et al. The Caribbean-Hispanic Alzheimer’s disease brain transcriptome reveals ancestry-specific disease mechanisms. Neurobiol Dis. Jan 2023;176:105938. doi:10.1016/j.nbd.2022.105938

60. Hu Y, Xu Y, Mao L, et al. Gene Expression Analysis Reveals Age and Ethnicity Signatures Between Young and Old Adults in Human PBMC. Front Aging. 2021;2:797040. doi:10.3389/fragi.2021.797040

61. Cao X, Ding L, Mersha TB. Development and validation of an RNA-seq-based transcriptomic risk score for asthma. Sci Rep. May 23 2022;12(1):8643. doi:10.1038/s41598-022-12199-0

62. Wang E, Lemos Duarte M, Rothman LE, Cai D, Zhang B. Non-coding RNAs in Alzheimer’s disease: perspectives from omics studies. Hum Mol Genet. Oct 20 2022;31(R1):R54–R61. doi:10.1093/hmg/ddac202

63. Marigorta UM, Denson LA, Hyams JS, et al. Transcriptional risk scores link GWAS to eQTLs and predict complications in Crohn’s disease. Nat Genet. Oct 2017;49(10):1517–1521. doi:10.1038/ng.3936

64. Liang Y, Pividori M, Manichaikul A, et al. Polygenic transcriptome risk scores (PTRS) can improve portability of polygenic risk scores across ancestries. Genome Biol. Jan 13 2022;23(1):23. doi:10.1186/s13059-021-02591-w

65. Scalco R, Saito N, Beckett L, et al. The neuropathological landscape of Hispanic and non-Hispanic White decedents with Alzheimer disease. Acta Neuropathol Commun. Jun 29 2023;11(1):105. doi:10.1186/s40478-023-01574-1

