## Supplementary for "MU-BRAIN: MUltiethnic Brain Rna-seq for Alzheimer INitiative"

Author affiliations:

1. Taub Institute for Research on Alzheimer’s Disease and the Aging Brain, College of Physicians and Surgeons, Columbia University. 630 West 168^th^ Street, New York, NY 10032, USA.
2. The Gertrude H. Sergievsky Center, College of Physicians and Surgeons, Columbia University. 630 West 168^th^ Street, New York, NY 10032, USA.
3. Department of Neurology, College of Physicians and Surgeons, Columbia University and the New York Presbyterian Hospital. 710 West 168^th^ Street, New York, NY 10032, USA.
4. Department of Pathology and Cell Biology, Columbia University, New York, NY, USA.
5. Department of Pathology and Laboratory Medicine, UC Davis Health Sacramento, CA 95817
6. Department of Neuroscience, Mayo Clinic Florida, Jacksonville, Florida, USA
7. Department of Neurosciences, University of California San Diego, La Jolla California, USA.
8. John P. Hussman Institute for Human Genomics, University of Miami Miller School of Medicine, Miami, FL, USA.
9. Department of Medical and Molecular Genetics, Indiana University, Indianapolis, Indiana, USA.

Correspondence to: Giuseppe Tosto, M.D., Ph.D.

G.H.Sergievsky Center, The Taub Institute for Research on Alzheimer's Disease and the Aging Brain

Department of Neurology, Columbia University Medical Center

630W 168^th^ street, room PH19-314

New York City, New York, 10032

**Running title**: Multiethnic transcriptome for AD

**Keywords:** Alzheimer’s Disease; Multi-ethnic; RNA-seq; Braak; Differential gene expression; Machine learning.

$${PTRS}_{i}=\beta_{1}g_{i1}+\beta_{2}g_{i2}+\cdots+\beta_{K}g_{iK}$$

(1)

where ***β*** denotes weight and ***g*** denotes Log_10_ of the normalized RNA-seq gene expression values. Four different regression logistic models were implemented for AUC calculations:

To refine our findings, we prioritized 30 DEG that were replicated in ROS/MAP *(***Table 3**), including: *ADAMTS2* (LFC=0.253, *P_adj_* =1.21$\times{10}^{-4}$)*, VGF* (LFC=-0.661, *P_adj_* =3.78$\times{10}^{-7}$), and *HSPB7* (LFC=0.445; *Padj*=3.78$\times{10}^{-7}$). **Supplemental Table 3** presents the top DEG identified by the transethnic metanalysis, including *ADAMTS2* (LFC_meta_=0.265, *P_adj_*_-meta_=1.00$\times{10}^{-16}$)*, VGF* (LFC_meta_ =-0.332, *P_adj_*_-meta_=2.24$\times{10}^{-7}$), and *HSPA6* (LFC_meta_ =-0.566; *P_adj_*_-meta_ =1.17$\times{10}^{-07}$).

### Data availability

Please contact the correspondence author for data related to this study.

### Funding

The project was partially funded by the National Institute on Aging (NIA) of the National Institutes of Health (NIH) under several Award Numbers: U01AG081817; R01AG082009; R01AG062517; R01AG67501; P30AG072972; P30AG062429; P30 AG062677; P01 AG003949; P30 AG062429 & AG066530; Florida Department of Health, Ed and Ethel Moore Alzheimer’s Disease Research Program (20A22, 8AZ06).

WHICAP: Data collection and sharing for this project was supported by the Washington Heights-Inwood Columbia Aging Project (WHICAP, PO1AG07232, R01AG037212, RF1AG054023, RF1AG066107 and R01AG072474) funded by the National Institute on Aging (NIA) and by the National Center for Advancing Translational Sciences, National Institutes of Health, through Grant Number UL1TR001873. This manuscript has been reviewed by WHICAP investigators for scientific content and consistency of data interpretation with previous WHICAP Study publications. We acknowledge the WHICAP study participants and the WHICAP research and support staff for their contributions to this study.

NIA-ADFBS: The NIA-LOAD study supported the collection of samples used in this study through National Institute on Aging (NIA) grants U24AG026395, R01AG041797 and U24AG056270. We thank contributors, including the Alzheimer’s disease Centers who collected samples used in this study, as well as patients and their families, whose help and participation made this work possible.

EFIGA: Data collection for this project was supported by the Genetic Studies of Alzheimer’s disease in Caribbean Hispanics (EFIGA) funded by the National Institute on Aging (NIA) and by the National Institutes of Health (NIH) (5R37AG015473, RF1AG015473, R56AG051876, R01AG067501). We acknowledge the EFIGA study participants and the EFIGA research and support staff for their contributions to this study.

### Competing interests

The authors report no competing interests.

### Supplementary material

Supplementary material is available at *Brain* online.

**REFERENCES.**

1.	Braak H, Braak E. Neuropathological stageing of Alzheimer-related changes. *Acta Neuropathol*. 1991;82(4):239-59. doi:10.1007/BF00308809
